# Aromatase Inhibitor Induced Musculoskeletal Inflammation is Observed Independent of Oophorectomy in a Novel Mouse Model

**DOI:** 10.1101/2022.06.22.497263

**Authors:** Nicholas A. Young, Jeffrey Hampton, Juhi Sharma, Kyle Jablonski, A. Courtney DeVries, Anna Bratasz, Lai-Chu Wu, Maryam Lustberg, Raquel Reinbolt, Wael N. Jarjour

**Affiliations:** Department of Internal Medicine, WVU Cancer Institute, West Virginia Clinical and Translational Science Institute, Morgantown, WV, 26506, USA; Division of Rheumatology and Immunology, WVU Cancer Institute, West Virginia Clinical and Translational Science Institute, Morgantown, WV, 26506, USA; Department of Neuroscience, Rockefeller Neuroscience Institute, Department of Medicine, Division of Hematology and Oncology, WVU Cancer Institute, West Virginia Clinical and Translational Science Institute, Morgantown, WV, 26506, USA; Small Animal Imaging Core, New Haven, CT, 06519, USA; Department of Biological Chemistry and Pharmacology, New Haven, CT, 06519, USA; Smilow Cancer Hospital/Yale Cancer Center, New Haven, CT, 06519, USA; The James Cancer Hospital, Columbus, OH, 43210, USA; The Ohio State University Wexner Medical Center, Columbus, OH, 43210, USA

**Keywords:** Breast Cancer, Hormone Receptor Positive Cancer, Estrogen, Aromatase Inhibitor, Aromatase Inhibitor Induced Arthralgia (AIIA)

## Abstract

**Background:** Aromatase Inhibitors (AIs) block physiological estrogen production in peripheral tissues and are used clinically to reduce disease recurrences and improve overall survival rates in hormone receptor-positive breast cancer patients. However, half of patients taking these drugs develop aromatase inhibitor induced arthralgia (AIIA), which is characterized by severe pain and inflammation in various joints and the surrounding musculoskeletal tissue. While the pathophysiology is not currently understood, it has been proposed to be associated with systemic estrogen deficiency resulting from AI treatment. Since AIIA leads to suspension of therapy in 20-30% of patients, reducing AIIA incidence may provide sustained AI treatment and enhance long-term survival.

**Objective:** In order to establish a better understanding of disease pathology and to create a platform that can be used to explore future interventional strategies, our objective in this study was to design a novel animal model of AIIA.

**Methods:** Female BALB/C-Tg(NFκB-RE-luc)-Xen mice, which have a firefly luciferase cDNA reporter transgene under the regulation of NFκB binding sites, were oophorectomized and treated with AI (letrozole) by daily subcutaneous injections for 5 weeks. Control groups included oophorectomized mice receiving vehicle injections and non-oophorectomized mice treated with AI. Knee joints and surrounding muscle tissue were imaged on the BioSpec 94/30 micro-MRI. The primary weight-bearing joint (hind limb) was examined histopathologically and NFκB activity was measured by bioluminescent imaging. Serum was collected for cytokine analysis. Additionally, healthy human PBMCs were treated with letrozole, estrogen, or both, and RNA sequencing was performed at 36 hrs.

**Results:** Bioluminescent imaging showed significantly enhanced NFκB activation with AI treatment in the hind limbs compared to controls receiving vehicle treatment. Moreover, analysis of knee joints and legs by MRI showed enhanced signal detection in the joint space and surrounding tissue following daily AI injections. Surprisingly, the enhanced MRI detection and NFκB activation was observed with AI treatment independent of the oophorectomy procedure. This indicates that the induction of musculoskeletal-directed inflammation by AI is not mediated by changes in physiological estrogen levels, which is contrary to proposed mechanisms of disease pathogenesis. Similarly, histopathological analysis showed tenosynovitis and musculoskeletal infiltrates in all mice receiving AI with or without oophorectomy. IHC analysis of the infiltrates demonstrated a predominantly macrophage-mediated inflammatory response with scattered CD4+ T cells. Additionally, serum cytokine levels of IL-2, IL-4, IL-6, and CXCL1 were significantly elevated in mice with AI treatment. RNA sequencing of human PBMCs after *in vitro* AI stimulation did not demonstrate an AI-specific gene expression pattern associated with immune system activation directly, suggesting that the pathogenesis of AIIA may be mediated through cells in other tissues *in vivo*.

**Conclusions:** Collectively, these data establish a novel mouse model of AIIA and identify an estrogen-independent stimulation of disease pathology via AI-mediated induction. This suggests that the pathogenesis of AIIA may not be mediated by estrogen deficiency, as previously hypothesized, and indicates that AI-induced inflammation may not be regulated directly through a pathogenic mechanism initially derived from circulating mononuclear cells. Future studies aim to characterize this inflammatory mechanism *in vivo* with a focus on other cells, including macrophages, synovial cells and chondrocytes, to provide insight into putative therapeutic strategies directed at mitigating disease pathology.

## INTRODUCTION

Aromatase Inhibitors (AIs) block physiological estrogen production in peripheral tissues and significantly improve overall survival rates by reducing disease recurrences in hormone receptor-positive breast cancer patients [1]. AI-induced arthralgias (AIIAs) are experienced by approximately 35-50% of women taking AIs and are characterized by joint stiffness, pain, tissue inflammation, and swelling [2-4]. These adverse events can significantly impact patient quality of life and contribute to early discontinuation and/or non-adherence to therapy [5] leading to termination in 10-20% of patients [3]. Moreover, recent reports suggest that poorer patient outcomes occur as a result of non-adherence/non-compliance with AI therapy [6]. and data indicate that AI treatment extension would offer more protection from tumor relapse.. In summary, AIIAs are a significant clinical obstacle to breast cancer treatment with AIs in the maintenance phase because of the high incidence of AIIA occurrence and the negative impact on outcome when treatment is suspended.

The diagnosis of AIIA is generally clinical, lacking standardized diagnostic criteria or confirmatory testing. Furthermore, there are no defined methods to objectively measure the effect of interventions designed to prevent or minimize toxicity. While the exact pathogenic mechanisms of AIIA remain to be definitively elucidated [4], estrogen deprivation directly mediated by AIs has been hypothesized and some preclinical studies have demonstrated estrogen to have a protective effect in arthritis and on the expression of pro-inflammatory genes [7-9]. Moreover, there is evidence that the pro-inflammatory cytokines IL-1, IL-6, and TNF-α are all increased in the first several years following menopause, which coincides with lower estrogen levels and a period during which joint symptoms are prevalent [10]. Others have postulated that estrogen may modulate spinal opioid anti-nociceptive activity to have a pain modulating effect [11]. Also, upregulation of inflammatory pathways and pro-inflammatory cytokines may be regulated by estrogen [12], especially in female predominant autoimmune diseases (reviewed in [13]). Collectively, a better understanding of the underlying molecular and pathogenic mechanisms of AIIA development and the role of estrogen specifically in this process may enhance efficacy of interventions aiming to prevent or treat this condition.

Translational animal models are a powerful tool to study the pathophysiology of human disease and to evaluate putative medical countermeasures. Although mouse models have been frequently used to study inflammation associated with rheumatologic disease, there are few well-developed preclinical models that could potentially be used to study the pathophysiology of AIIAs. Letrozole is the most potent oral AI used in breast cancer patients [8] and was used in this study to determine whether inflammation can indeed be detected following treatment. In order to recapitulate a post-menopausal state, an oophorectomy procedure using previously established animal models was performed on female mice [14]. This well-established model has demonstrated more than a 30% reduction in systemic estrogen levels approximately two weeks after the procedure [15]. Following daily treatment with letrozole, our results demonstrate successful detection of AI-induced inflammatory responses and establish a novel animal model of AIIA. Interestingly, AI-induced inflammation was observed irrespective of the oophorectomy procedure, which indicates that AIIA pathogenesis is independent of systemic physiological estrogen levels.

## METHODS

### AIIA induction and sample size determination

BALB/C-Tg(NFκB-RE-luc)-Xen mice (1.5 - 6 mo of age) carrying a transgene containing six NFκB responsive elements and a modified firefly luciferase cDNA were purchased from Caliper Life Science (Hopkinton, MA). Female NFκB-RE-luc mice were oophorectomized (N = 15) and, after a 4-weeks recovery period, treated with AI (letrozole; Sigma-Aldrich, St. Louis, MO) by daily subcutaneous injections (10 µg/mouse/day) for 5 weeks. Control groups included oophorectomized mice (OVX) receiving vehicle (0.3% hydroxypropyl cellulose; HPC) injections (N = 10), and non-oophorectomized mice (sham surgery performed) treated with AI (N = 15). An experimental summary of the AIIA animal model protocol is shown in **Figure 1**. Mouse maintenance and protocols were approved by the Institutional Animal Care and Use Committee at The Ohio State University Wexner Medical Center (OSUWMC). The animal facility was maintained at 22-23°C and 30-50% relative humidity with a 12-hour light/dark cycle; chow and water were supplied ad libitum. The sample sizes were based on calculations and data from previous studies with type one error of 0.05 and power of 0.8 [9]. Specifically, the power calculation was performed with regard to the effect on letrozole-mediated treatment of status epilepticus. We anticipated that we would be able to reject the null hypothesis, as long as the true difference between the groups is 0.8 or greater on the probability scale.

**Figure 1.**
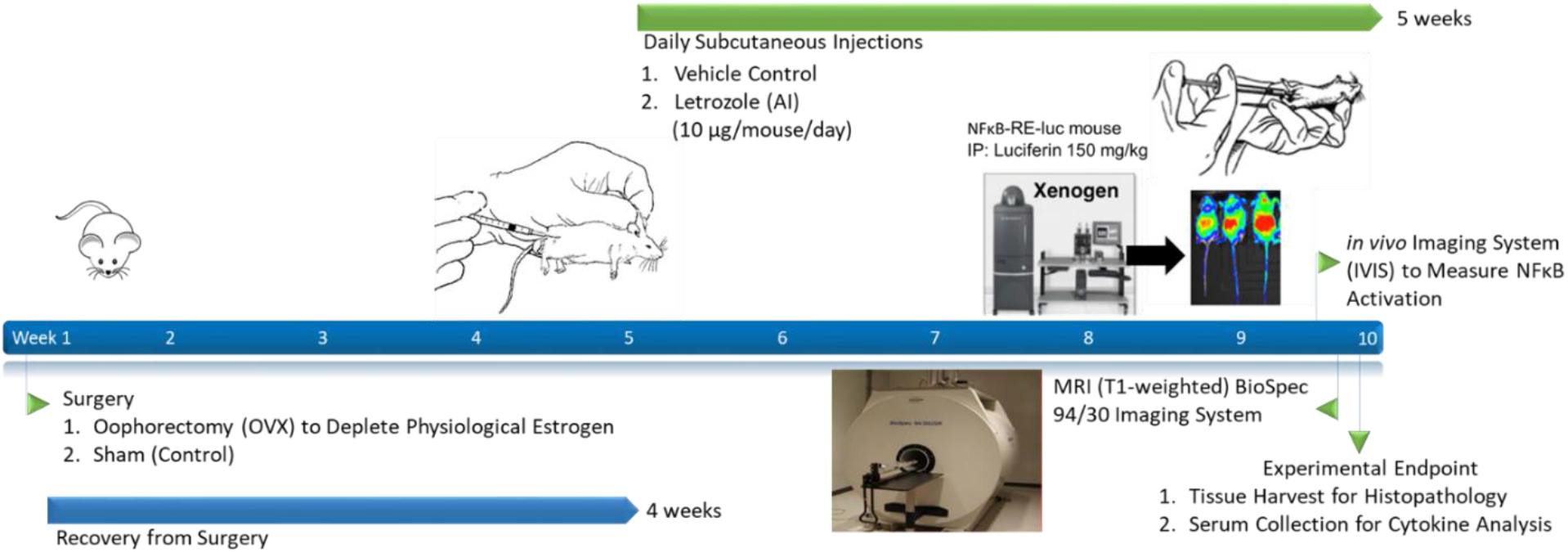
Schematic of experimental outline and sample collection establishing a novel murine model of AIIA.

### Tissue collection

Tissues were resected from each mouse and flash frozen in liquid nitrogen-cooled isopentane, followed by cryosectioning, or immediately fixed by immersion in neutral buffered 10% formalin and then processed into paraffin. Serial histological sections were stained with H&E (Leica Microsystems Inc., Buffalo Grove, IL) following the manufacturer’s protocol, or labeled by immunohistochemistry (IHC) as detailed previously [16] and per established protocols at HistoWiz (Brooklyn, NY). Histological slides were scanned for downstream digital/quantitative image analysis.

### Histopathology

H&E-stained paraffin sections of tissue were subjected to blinded histopathological analysis using the 10x objective of a bright-field light microscope with predefined criteria [16, 17]. Scanned slide images were analyzed by Aperio ImageScope digital analysis software (v9.1) as detailed previously to determine positive staining and lymphocyte localization by immunohistochemistry [17, 18]. Specifically, Aperio’s positive pixel count algorithm was run to quantify the extent of positive staining and lymphocyte localization using calibrated hue, saturation, and intensity values following previously described methods of computer-assisted image analysis [19]. To define a mean positive pixel intensity value, 10 measurements of identical total surface area of tissue were quantitated. All digital analysis was confirmed by manual slide interpretations using the 10x objective of a bright-field light microscope.

### Cytokine measurement

Serum was prepared from whole blood collected via subclavian artery at the experimental endpoint. Serum samples were analyzed by enzyme-linked immunosorbent assay (ELISA) using the V-PLEX Pro-inflammatory Panel 1 mouse kit (Meso Scale Diagnostics, Rockville, MD) according to the manufacture’s protocol and data was analyzed using Microsoft Excel (v2204), as previously described [16]. The limit of detection (LOD) using this kit with each analyte is: IL-2 (0.12 pg/mL), IL-6 (0.12 pg/mL), IFN-γ (0.05 pg/mL), KC/GRO (CXCL1, 0.05 pg/mL), IL-4 (0.04 pg/mL), and IL-5 (0.07 pg/mL). All reported results were above the LOD of the assay.

### Bioluminescent Imaging with IVIS

All IVIS imaging and analysis was performed as described previously [3]. In brief, BALB/C-Tg(NFκB-RE-luc)-Xen mice were given 150 mg/kg luciferin (15 mg/mL in PBS pH 7; unadjusted) through intraperitoneal (IP) injection and bioluminescent signals were captured using the IVIS 200 system (Caliper Life Sciences). Data were quantified and analyzed using IVIS Living Image^®^ software (v3.2).

### MRI

The BioSpec 94/30 Imaging System with a 9.4T horizontal bore magnet that operates at 400 MHz using ParaVision(tm) 5.1 software. MRI scans were acquired with a 72 mm volume coil (transceiver) with following acquisition parameters: MSME (multi spin multi echo) method, echo time = 6.834ms (minimum), 30 echos (6.834-136.68), repetition time = 2000ms, averages = 2, resolution: 150*150 micron (FOV vary, but resolution constant), and slice thickness of 1 mm.

### Letrozole stimulation of human PBMCs

PBMCs freshly isolated from healthy pre-menopausal females were cultured in hormone free conditions as previously described [20] using RPMI 1640 (Life Technologies, Grand Island, NY) and 5% charcoal stripped fetal bovine serum (FBS; Life Technologies). Participation was in accordance with an approved Institutional Review Board protocol at OSUWMC. PBMCs were treated with 10 nM of 17β-estradiol (E2; Sigma) and/or 10 nM of letrozole (Sigma) and were collected at 36 hrs according to previously established protocols [12].

### Statistics

Statistical analysis of cytokine detection and IVIS measurements for three groups collectively was performed by one-way ANOVA. Digital pixilation of IHC staining was analyzed by two-way ANOVA to estimate mean quantitative variable changes according to the levels of both experimental groups (letrozole oophorectomy and letrozole sham) relative to each IHC stain. All ANOVA analyses were followed by Tukey’s post hoc test for multiple comparisons and numerical datasets were expressed as mean values with standard error of the mean indicated (SEM) using Graph Pad Prism software (v8.3.0). Data was considered statistically significant if p ≤ 0.05.

## RESULTS

### Letrozole-induced inflammation detection by MRI and IVIS in murine hind limbs

After 5 weeks of daily subcutaneous letrozole injections, knee joints and surrounding hind limb tissues of mice were imaged on the BioSpec 94/30 micro-MRI with gadolinium enhancement under anesthesia. Analysis of knee joints and proximal anatomy by MRI showed enhanced signal detection in the joint space and surrounding tissue following letrozole treatment in oophorectomized NFκB-RE-luc mice (**Figure 2**). Surprisingly, enhanced MRI detection was also demonstrated in non-oophorectomized (sham surgery) mice that were treated with letrozole. Although the MRI signal detection was lower without the oophorectomy procedure, letrozole did elevate responses relative to untreated controls (daily vehicle injections following an oophorectomy procedure), which indicates that AIIA pathophysiology may be independent of physiological levels of estrogen.

**Figure 2.**
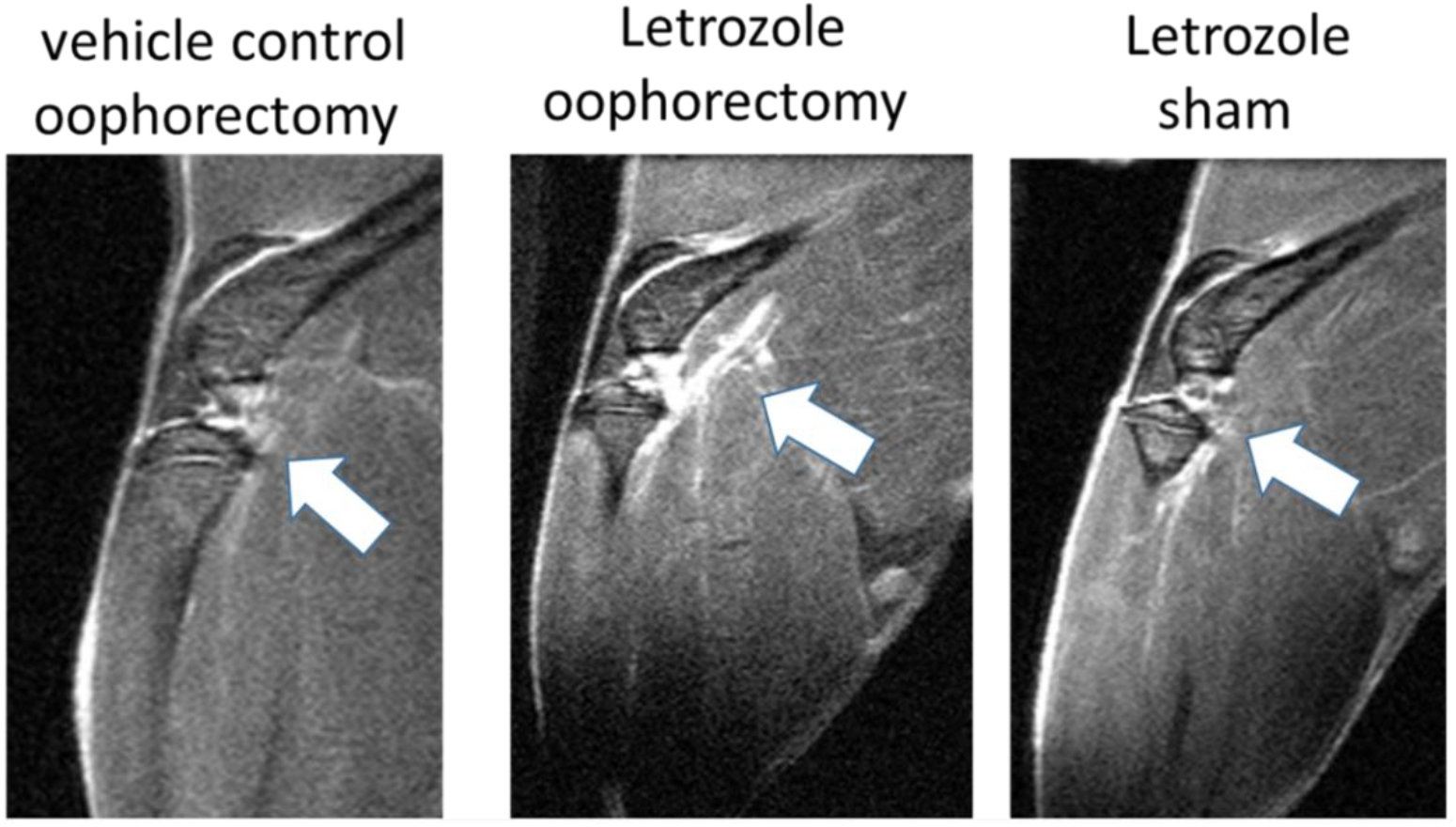
Enhanced MRI signals detected in the legs of mice receiving aromatase inhibitor treatment. Female NFkB-RE-luc mice were injected subcutaneously with the aromatase inhibitor letrozole (10µg/mouse/day) or vehicle control (0.3% hydroxypropyl cellulose) following oophorectomy or control surgery (sham). MRIs were conducted on live mice after 5 weeks of treatment. Enhanced signals (white arrows) are observed in knees and surrounding muscle tissue by MRI (T1-weighted) with letrozole treatment and these responses were detected in NFkB-RE-luc mice irrespective of the oophorectomy procedure.

The transcription factor NFκB serves as a pivotal mediator of inflammatory responses. In order to measure NFκB activation that might associate with aromatase inhibitor treatment, bioluminescent imaging was performed on NFκB-RE-luc mice after 5 weeks of letrozole treatment using luciferin and the *in vivo* imaging system (IVIS). Digital quantitation of emitted photons from the hind limbs showed significantly enhanced NFκB activation with letrozole treatment compared to oophorectomized controls receiving vehicle injections (**Figure 3**). Specifically, letrozole-mediated NFκB activation increased 45% (p = 0.04) with an oophorectomy and 65% (p = 0.0006) in mice with sham surgeries. In concordance with the MRI data above, the activation of NFκB was observed in both mouse groups with and without oophorectomy, which suggests that the induction of letrozole-induced inflammatory responses is not dependent on estrogen levels.

**Figure 3.**
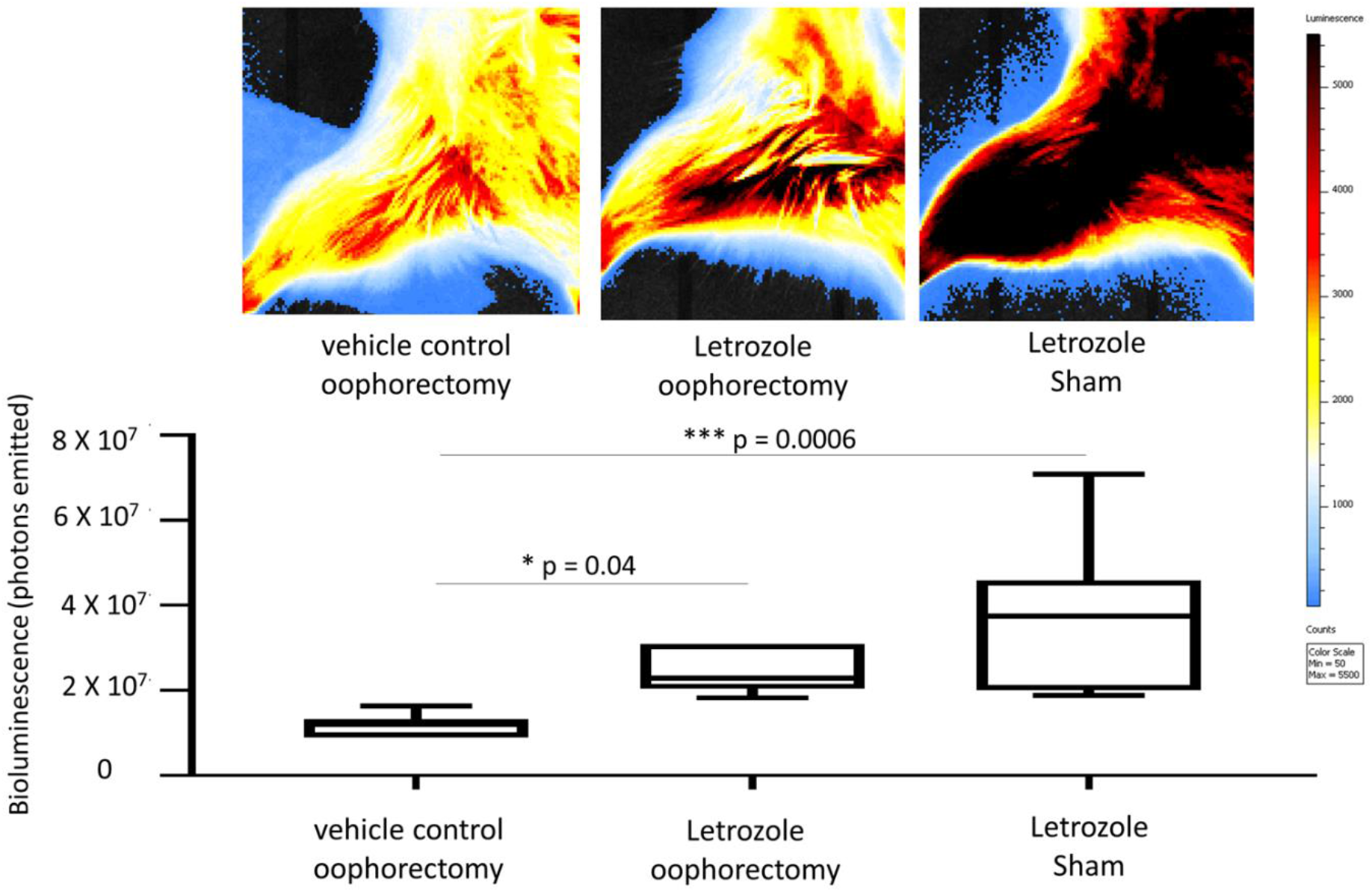
Bioluminescent imaging of NFκB activity shows aromatase inhibitor-induced upregulation with and without oophorectomy. Female NFkB-RE-luc mice were treated with letrozole following an oophorectomy procedure. Controls included sham surgery and vehicle control treatment (0.3% hydroxypropylcellulose; HPC). NFκB activity was measured on the *in vivo* imaging system (IVIS) approximately at the 5 week timepoint. IVIS (top) and quantitation of bioluminescent signals (bottom) demonstrate induction of NFκB activity both with and without oophorectomy relative to control mice. Values are the mean ± SEM with indicated p values calculated via one-way ANOVA with Tukey’s corrections for multiple comparisons. *All values of p ≤ 0.05 considered statistically significant.

### Histopathological analysis shows that letrozole increase macrophage infiltration

At the experimental endpoint, the hind limbs of mice were collected for histopathological analysis. H&E staining of tissue sections demonstrated mild tenosynovitis and inflammatory muscle tissue infiltrates (**Figure 4**) in both groups of mice treated with letrozole with and without oophorectomy. To characterize the immune cell subtypes present in these infiltrates, IHC was performed to detect CD4+ T cells, CD8+ T cells, B220+ B cells, F 4/80+ macrophages, and Ly6G+ neutrophils of tendons and ligaments (**Figure 5**) as well as other musculoskeletal tissue (**Figure 6**). The results showed that the infiltrates in those tissues were consisted largely of macrophages. Digital image analysis via two-way ANOVA was used to estimate how the mean quantitative values changed according to the levels of categorical variables. Here, the categories compared statistically were immune cell subtype (i.e., IHC staining category). The data demonstrate that macrophage detection relative to the other immune cell subtypes was greatest in areas of tenosynovitis (p < 0.0001), but also significantly elevated in muscle tissue (p < 0.0002). Additionally, CD4+ T cells were also detected to a lesser extent in histopathological analysis of tendons/ligaments and muscle tissue; digital quantification of CD4 IHC was statistically significant in tenosynovitis infiltrates with letrozole stimulation when compared to the other immune cell subtypes (p = 0.05).

**Figure 4.**
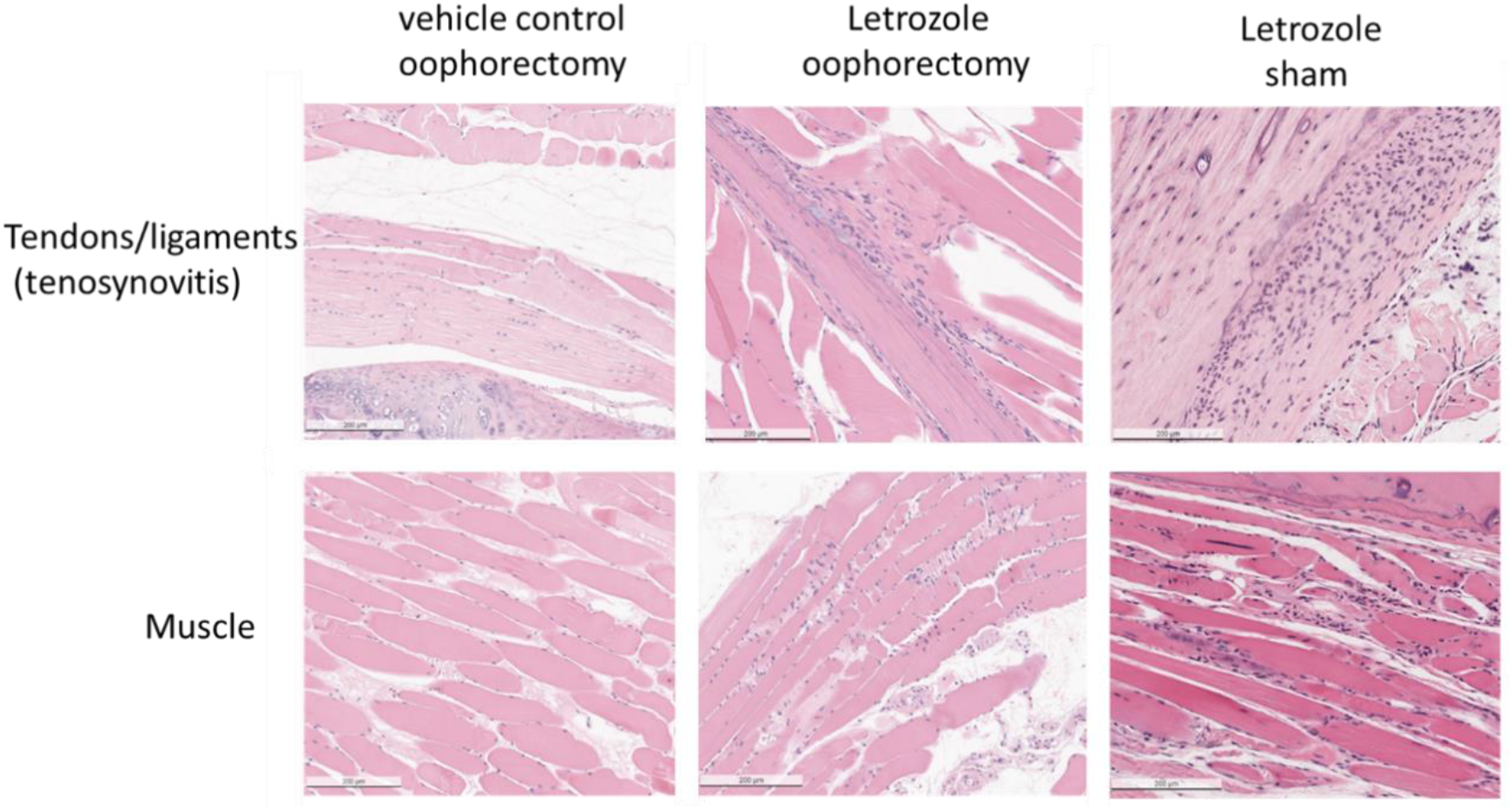
Histopathological analysis of mice receiving aromatase inhibitor treatment indicates tenosynovitis and myositis. H&E staining of tissue sections taken from female NFκB-RE-luc mice after 5-weeks of subcutaneous letrozole treatment. Muscle tissue sections were dissected from legs and processed for H&E staining and histopathological analysis. Cellular infiltrates are present in tendons and muscles with treatment of the aromatase inhibitor letrozole independent of oophorectomy or sham surgery.

**Figure 5.**
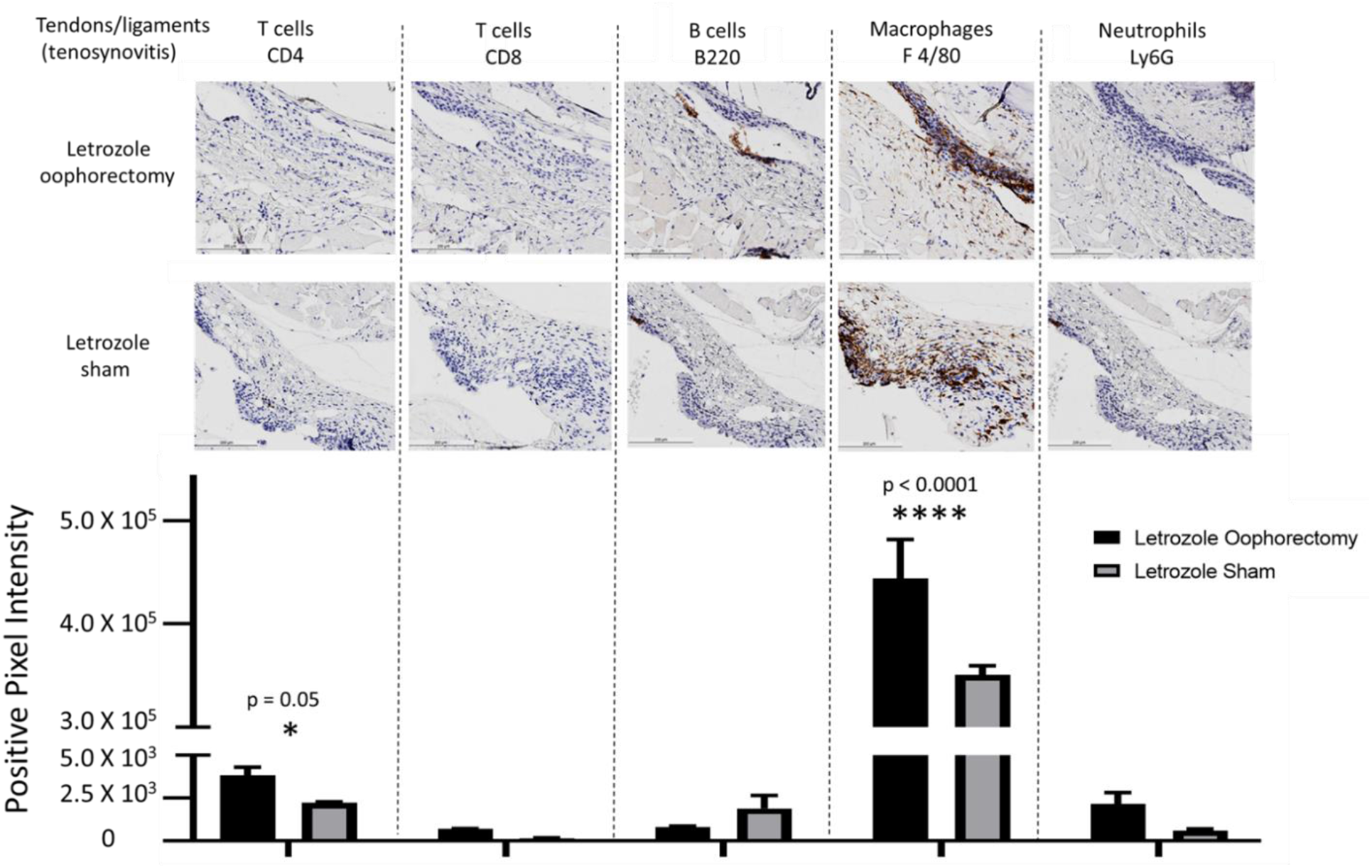
IHC for immune cell subtypes in tenosynovitis infiltrates shows a macrophage-mediated inflammatory response induced by aromatase inhibitor treatment. Following oophorectomy or control (sham) surgery and induction of AIIA with daily letrozole injections, mouse legs were collected and processed for histpathological assessment as described in the materials and methods. To evaluate immune cell subtypes in the tendons, IHC staining of CD4+ T cells, CD8+ T cells, B220+ B cells, F 4/80+ macrophages, and Ly6G+ neutrophils was performed. Data were analyzed by two-way ANOVA for group-wise comparisons relative to each IHC stain. Values are the mean ± SEM with indicated p values determined following Tukey’s corrections for multiple comparisons to measure the statistical change between IHC cell stain/subtype. *All values of p ≤ 0.05 considered statistically significant compared to all other stains individually.

**Figure 6.**
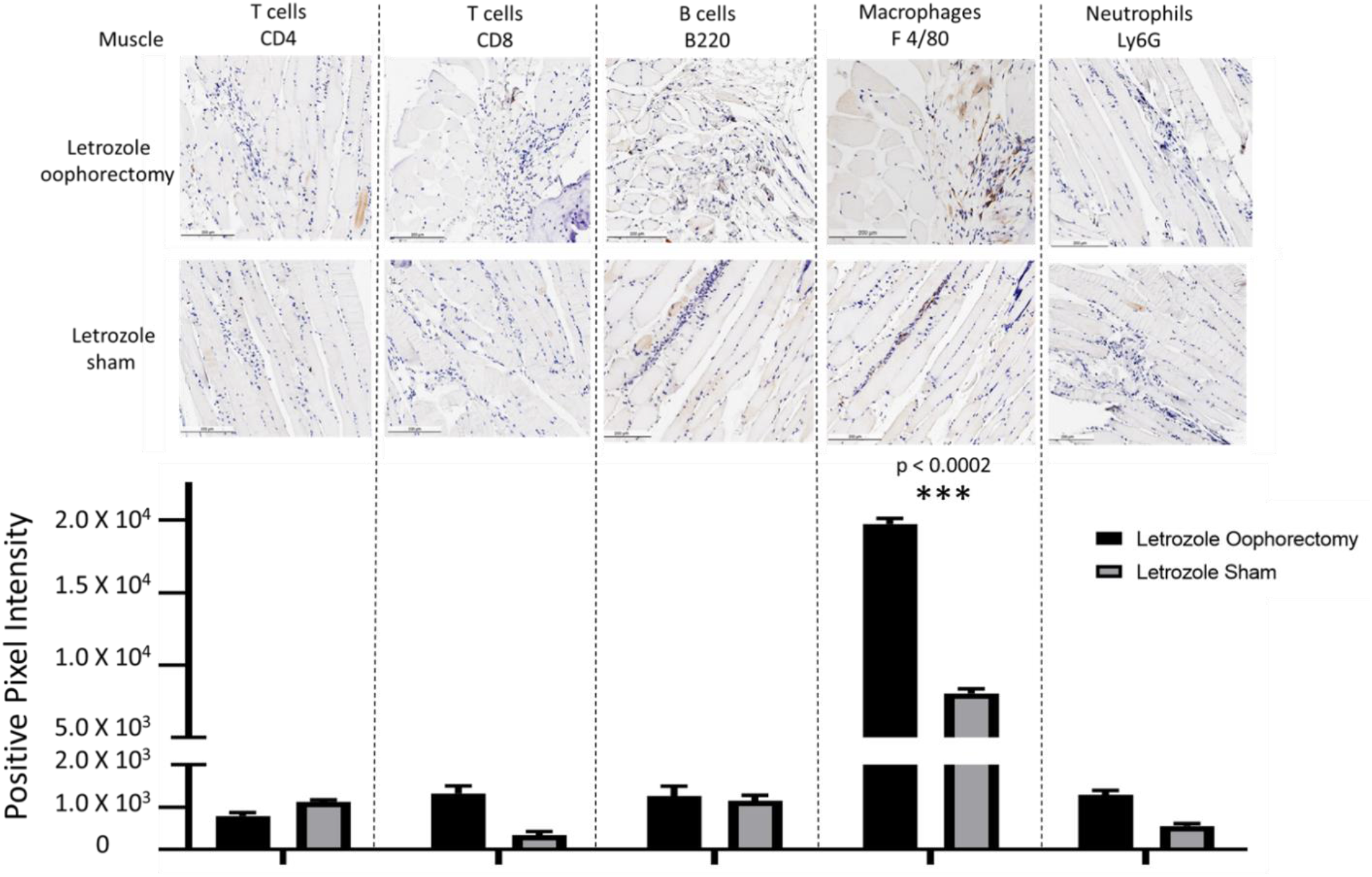
Macrophage predominant inflammation of muscle tissue detected with AIIA induced by aromatase inhibitor treatment. As outlined in the materials and methods, AIIA was induced in mice by administration of daily letrozole injections subsequent to oophorectomy or control (sham) surgery. Histopathology of muscle tissue was evaluated by IHC staining of CD4+ T cells, CD8+ T cells, B220+ B cells, F 4/80+ macrophages, and Ly6G+ neutrophils to characterize immune cell subtypes. Data analysis was performed using two-way ANOVA for group-wise comparisons relative to each IHC stain. Two-way ANOVA with Tukey’s correction for multiple comparisons was performed to evaluate the statistical changes between IHC cell stain/subtype. Values shown are the mean ± SEM with indicated p values relative to other immune cell subtypes. *All values of p ≤ 0.05 considered statistically significant relative to each stain individually.

### AIIA mouse model shows induction of pro-inflammatory cytokines

To measure systemic inflammation biomarkers, serum was prepared from mice at the experimental endpoint and analyzed by electrochemiluminescent ELISA. Pro-inflammatory cytokine levels of IL-2, IL-4, IL-6, and KC/GRO (CXCL1) were significantly elevated with letrozole treatment when compared to untreated, oophorectomized control mice irrespective of the oophorectomy procedure (**Figure 7**). Relative to oophorectomized mice receiving vehicle control injections, letrozole induced systemic IL-4 (p = 0.04), IL-6 (p = 0.03), IL-2 (p = 0.02), and KC/GRO (CXCL1) (p < 0.0001) following the oophorectomy procedure. Despite a lack of physiological estrogen depletion following sham surgery, letrozole significantly enhanced serum expression of IL-4 (p = 0.04), IL-6 (p = 0.0002), IL-2 (p = 0.002), and KC/GRO (CXCL1) (p < 0.03). These data further support the hypothesis that letrozole is inducing a pro-inflammatory response in this murine model of AIIA independent of estrogen levels.

**Figure 7.**
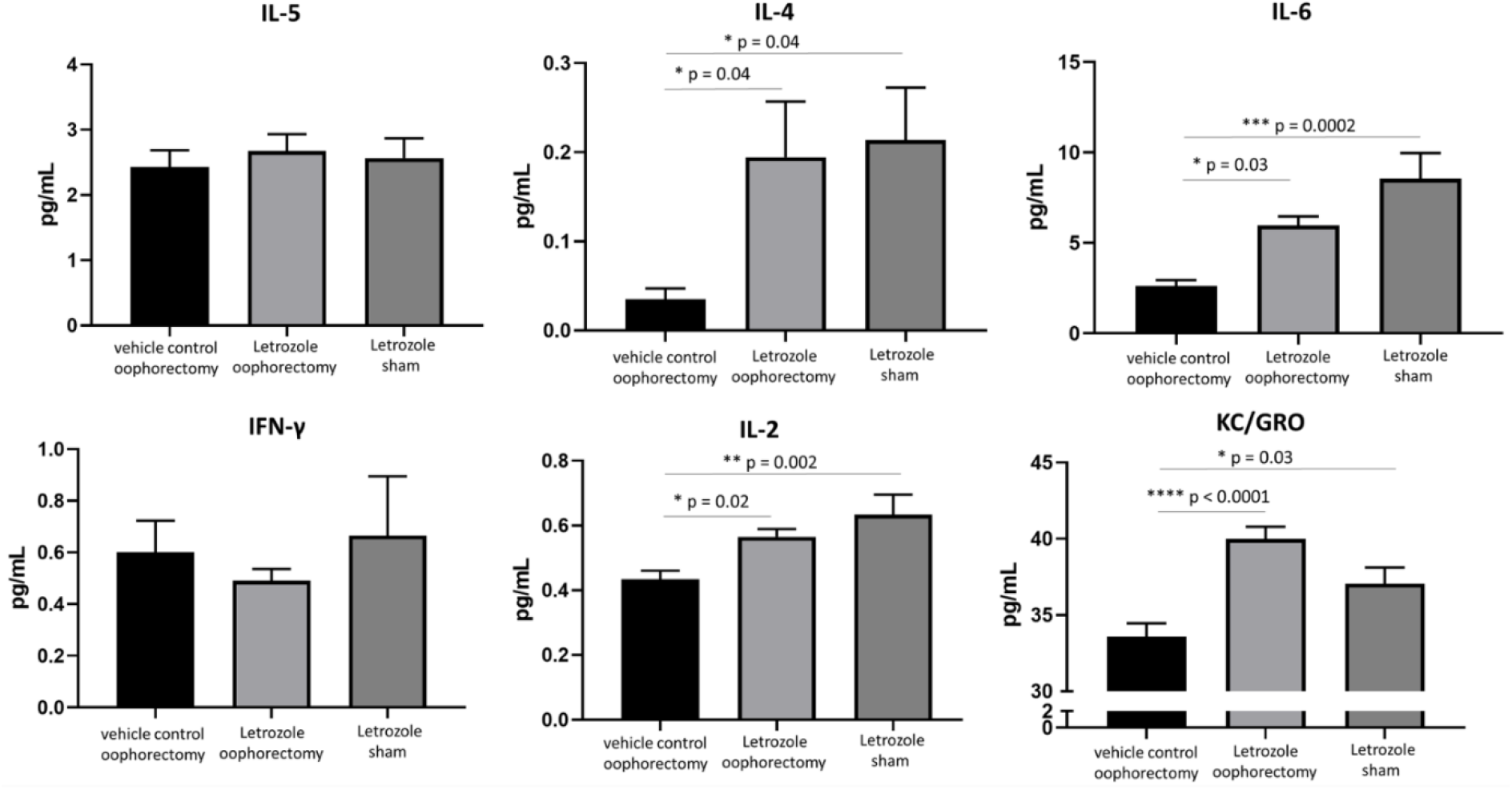
Aromatase inhibitors induce pro-inflammatory cytokine levels in the serum of NFκB-RE-luc mice independent of physiological estrogen depletion. Female NFκB-RE-luc mice underwent oophorectomy or control (sham) surgery and were treated with the aromatase inhibitor letrozole for comparison to negative (vehicle) controls. Cytokine levels from serum were measured by electrochemiluminescence detection using a pro-inflammatory panel. Values are the mean ± SEM with indicated p values calculated from one-way ANOVA with Tukey’s corrections for multiple comparisons. *All values of p ≤ 0.05 considered statistically significant.

### Letrozole treatment of human PBMCs in vitro does not produce an observable genomic effect

To explore the genomic influence of letrozole in human PBMCs and to provide insight into the molecular mechanisms of inflammation in AIIA, healthy human PBMCs from 5 pre-menopausal females were isolated and then stimulated under 4 conditions: i) untreated, ii) estrogen, iii) letrozole, and iv) estrogen + letrozole. Cell pellets were collected at 36 hrs and RNA sequencing (RNAseq) for mRNA and (micro) miRNA transcriptional targets was performed. Targets with low counts were removed via edgeR::filterByExpr, which uses embedded filtering for sequencing depth and experimental design considerations. After filtering, 24,511 genomic targets were included in the mRNA analysis and 1,599 were included in the miRNA analysis; this correlates to 85% and 92% of the total, respectively. Normalization was done through the trimmed mean of M-values (TMM) method and indicated that the filtering was adequate for downstream comparative evaluation. Using these data, results were normalized to untreated values for each patient individually to account for patient-to-patient variability in the datasets. To evaluate genome-wide effects of these treatments, heatmaps were created for mRNA and miRNA targets with hierarchical clustering using Pearson’s correlation distance measure and Ward’s clustering criterion with squared dissimilarities. Rows were scaled so that sample locations could be compared across a gene and the number of genes was reduced to >10,000 for mRNA by further filtering genes that had the lowest standard deviations. The results for both mRNA and miRNA did not demonstrate any strong or biologically meaningful clustering (**Figure 8**), which indicates that induction of AIIA is unlikely mediated by a direct effect on the regulation of PBMC gene expression.

## DISCUSSION

AIs, including letrozole, are approved prophylactic, long-term treatments for hormone receptor positive breast cancer patients [1]. Although this adjuvant treatment strategy has demonstrated efficacy in reducing disease relapse via suppression of physiological estrogen levels, one complicating immune related adverse event is AIIA. AIIA is characterized clinically by joint pain, tenosynovitis, and musculoskeletal inflammation [2-4] with the proposed pathogenesis resulting presumably from estrogen deprivation [9-11]. Unfortunately, approximately half of patients receiving AI therapy develop AIIA, which results in treatment discontinuation in up to 20% of this cohort [3]. Since suspension of AI therapy is associated with increased risk of relapse [6], prevention or proactive management of AIIA could lead to better patient outcomes via prolonged adherence to treatment. With early breast cancer detection leading to better overall patient outcomes and AI treatment recommendations extending from five to ten years [7], more patients will inevitably develop AIIA. To establish an animal model of AIIA, we treated mice with letrozole and studied the responses both with and without oophorectomy. The results from this study establish a novel animal model of AIIA that can be leveraged in future work to evaluate novel interventions and diagnostics to prevent or ameliorate AIIAs, which may ultimately translate into improved clinical outcomes.

**Figure 8.**
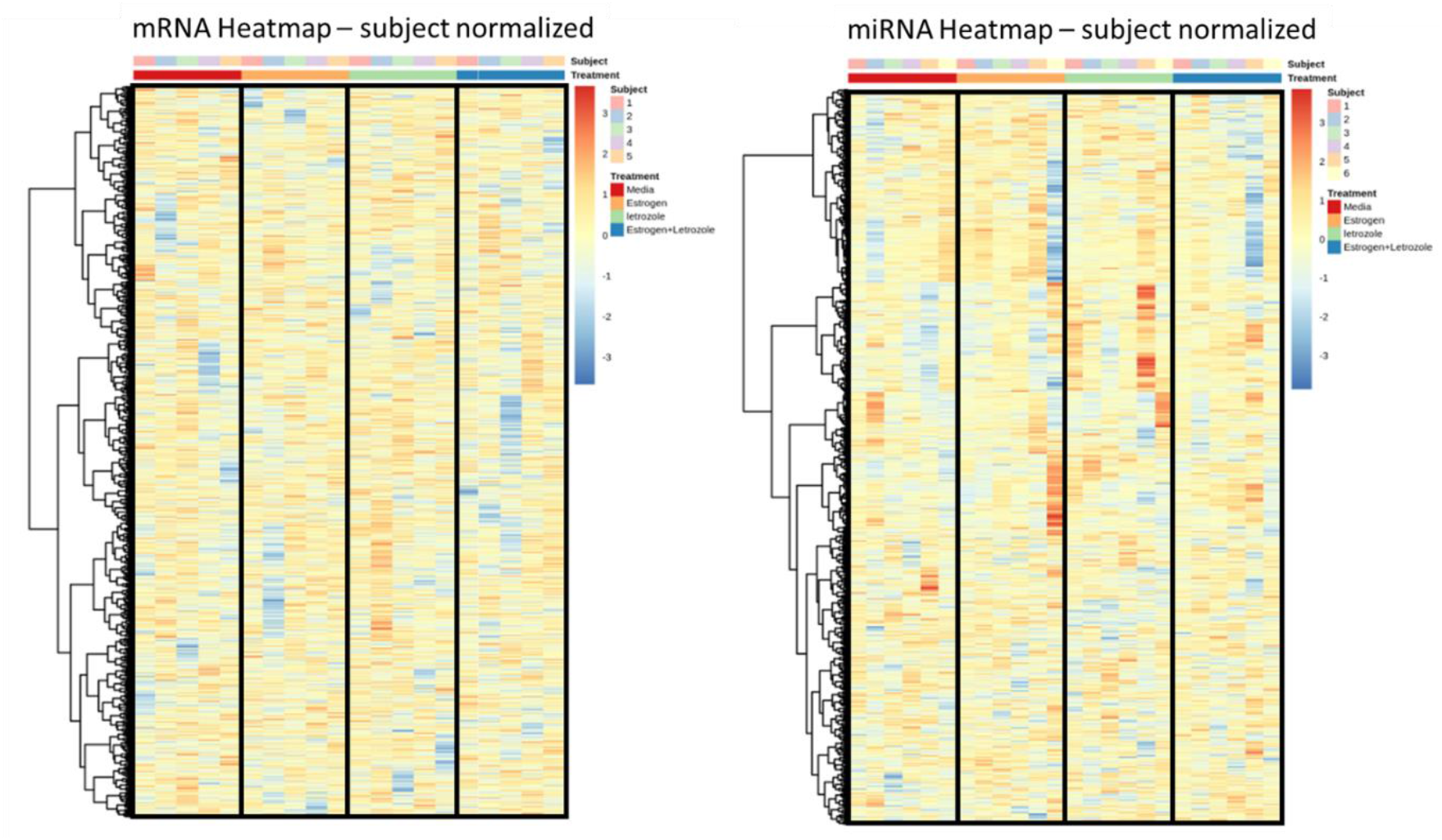
Genome-wide expression analysis of human PBMCs does not demonstrate an aromatase inhibitor-mediated response. Freshly isolated human PBMCs were stimulated *in vitro* with estrogen, letrozole, or a combination of both for comparison relative to untreated control samples. RNA sequencing was preformed to evaluate overall expression levels of mRNA and miRNA targets with each treatment sample. Data from each subject was normalized to control values and relative expression levels are presented as heatmaps to evaluate genome-wide treatment effects on mRNA (left) and miRNA (right) transcripts.

Our results demonstrated macrophage predominated infiltrates into both the tendons and surrounding musculoskeletal tissue and significantly induced levels of IL-4, IL-2, and IL-6 as well as chemokine KC/GRO (CXCL1) with daily letrozole injections. To the best of our knowledge, this is the first study to characterize AIIA-associated infiltrates histopathologically; therefore, these data may help inform putative AIIA prophylaxis options to evaluate in future studies. Since the inflammatory response consists mostly of macrophages, treatments directed at this specific immune cell subtype should be prioritized. Accordingly, in our previous studies evaluating the anti-inflammatory effects of nano-emulsified curcumin, we demonstrated a reduction in NFκB activation, macrophage-specific chemoattractant MCP-1 secretion, *in vitro* macrophage migration using both primary cells and cell lines, and macrophage recruitment in a mouse model of peritonitis [21]. Furthermore, histopathological analysis of a murine model of gout that we developed showed that the infiltrates in the tibio-tarsal joints (ankle) consisted largely of macrophages and neutrophils [22]; subsequent intervention with daily tart cherry juice concentrate significantly reduced macrophage-mediated inflammatory responses, as shown by histopathological analysis [23]. Additionally, therapeutic strategies leveraging selective macrophage phagocytosis of lipid particles loaded with immunomodulatory biocargo should also be considered [24] (similar to our strategy used in curcumin-containing lipid nanoparticles [21]). In addition to these potential interventions, the use of disease-modifying antirheumatic drugs (DMARDs) that have demonstrated the capacity to drive macrophage polarization from the pro-inflammatory M1 state to the wound healing M2 state may also be effective treatments for AIIA [25]. Specifically, Abatacept (small molecule inhibitor that blocks T cell activation by binding to CD80 or CD86 extracellularly to prevent interaction with co-stimulatory receptor CD28), Etanercept (soluble TNF receptor that competitively binds TNF-α and TNF-β), Infliximab (mAb that binds to TNF-α), Rituximab (mAb that binds to CD20 and inhibits B cell activation), and Tocilizumab (anti-IL-6 mAb) are all DMARDs to consider in future studies evaluating the efficacy of preventing or treating AIIA. Detection of enhanced systemic pro-inflammatory cytokine/chemokine expression with letrozole treatment indicates that these pro-inflammatory mediators are stimulating the observed macrophage-mediated inflammatory responses. Considering that IL-2 and IL-6 are well-characterized pro-inflammatory Th1-mediated cytokines [26] and KC/GRO (CXCL1) is associated with M1 polarization [27], these immunomodulators could also be selectively targeted to suppress AIIA pathogenesis. While overexpression of the Th2/M2-associated, wound healing cytokine IL-4 may be a compensatory mechanism that has been previously observed in inflammatory disease [28], this hypothesis will require additional investigation to elucidate more definitively. Collectively, the results from this study indicate that macrophage-mediated therapeutic targets associated with M1 polarization and inflammatory disease pathogenesis should be prioritized in future strategies to ameliorate AIIA. While letrozole treatment did result in both tenosynovitis and musculoskeletal inflammation independent of physiological estrogen levels, *in vitro* treatment of human PBMCs did not result in a measurable effect over gene expression. Consequently, these results suggest that the pathogenesis of letrozole-induced genetic responses may be derived from cell types other than PBMCs. In concordance, the interaction of synovial fibroblasts with tissue resident macrophages has been shown to be associated with synovitis in autoimmune disease pathogenesis [29]. In this model of rheumatoid arthritis, fibroblast secretion of IL-6, CXCL12, and CCL2 in the synovial microenvironment as well as expression of an IFN-γ gene signature stimulates the macrophage-mediated inflammatory response. Moreover, chondrocytes have a well-established role in the inflammatory disease pathogenesis of arthralgias and should also be considered in future *in vivo* studies evaluating the mechanism of AIIA induction. Chondrocytes can secrete IL-1β into the synovium to stimulate NFκB expression in immune cells and MMP-13, which degrades the extracellular matrix in the synovial compartment [30]. Consequently, activation of macrophages in the synovium by chondrocytes and/or synovial fibroblasts may be the impetus of a broader inflammatory response ultimately leading to tenosynovitis and musculoskeletal inflammation. Future studies using our murine model will characterize the roles of both chondrocytes and synovial fibroblasts as well as tissue resident macrophages in letrozole-induced inflammatory responses.

Interestingly, our data indicate that letrozole-induced inflammatory responses occur independent of physiological estrogen levels, as mice both with and without oophorectomy produced similar results in this study. This is particularly important because current prevailing hypotheses speculate that AIIA pathogenesis is mediated by estrogen suppression [9-11]. However, while AI treatment does suppress physiological estrogen, reduction systemically is not consistent from patient-to-patient and is largely inconsequential in clinically obese patients with extended treatment [31]. These data suggest that non-obese patients with statistically lower levels of estrogen following AI treatment should have a higher incidence of AIIA. On the contrary, obese patients are actually more likely to develop AIIA than their non-obese counterparts, indicating that AI-mediated induction of AIIA may indeed be estrogen-independent [32]. Similarly, estrogen replacement therapy has been shown to be efficacious in post-menopausal women with arthralgia not resulting from joint replacement due to hip fracture, according to the Women’s Health Initiative [33]. Considering these observations from human studies along with the results from our experiments, the prevailing estrogen-dependent hypothesis explaining the pathogenesis of AIIA should be further investigated and reconsidered in the context of AI-mediated effects independently.

These results establish a novel mouse model of AIIA; thus, our future studies with this model will be directed towards the further characterization of this inflammatory mechanism to provide insight into potential therapeutic strategies directed at mitigating this adverse inflammatory burden. Additionally, analysis of PBMCs and serum from breast cancer patients longitudinally with AI therapy could identify diagnostic biomarkers, relevant molecular pathways of disease pathogenesis, and potential therapeutic targets to correlate with murine samples from this AIIA model.

## ACKNOWLEGEMENTS

We would like to extend our gratitude to the Clinical Research Center and the Center for Clinical and Translational Science (CCTS) at Ohio State University’s Wexner Medical Center as well as the volunteers that participated in this study.

## COMPETITING INTERESTS

None declared.

## DATA SHARING STATEMENT

All of the data collected during this study, the data analysis plan, and the treatment protocol are on-file for sharing immediately following publication to anyone who wishes to access the data for any purpose with no end date. Proposals should be directed to. To gain access, requestors will need to sign a data access agreement.

## PROVENANCE AND PEER REVIEW

Not commissioned; externally peer reviewed.

## PATIENT CONSENT FOR PUBLICATION

Patient consent was in accordance with an approved Institutional Review Board protocol at OSUWMC.

## AUTHOR CONTRIBUTIONS

Conceived and designed the experiments of the study: (NY, ACD, ML, RR, WJ). Data collection and analysis: (NY, JH, JS, KJ, AB). Performed experiments: (NY, JH, JS, KJ, AB). Edited manuscript: (NY, JS, KJ, ACD, AB, LW, ML, RR, WJ). Statistical assessments: (NY, JH, JS, KJ). Wrote manuscript: (NY, ML, RR, WJ). Contributed reagents/materials/analysis tools: (ACD, AB, LW, WJ). Made substantial, direct and intellectual contribution to the work, and approved it for publication: (NY, JH, JS, KJ, ACD, AB, LW, ML, RR, WJ).

